# SS-31 Reverses Mitochondrial Fragmentation in Fibroblasts from Patients with DCMA, a Mitochondrial Cardiomyopathy

**DOI:** 10.1101/672857

**Authors:** Pranav Machiraju, Xuemei Wang, Rasha Sabouny, Joshua Huang, Tian Zhao, Fatima Iqbal, Melissa King, Dimple Prasher, Arijit Lodha, Amir Ravandi, Bob Argiropoulos, David Sinasac, Aneal Khan, Timothy Shutt, Steven C. Greenway

## Abstract

**Objectives:** We used patient dermal fibroblasts to characterize the mitochondrial abnormalities associated with the dilated cardiomyopathy with ataxia syndrome (DCMA) and to study the effect of the mitochondrially-targeted peptide SS-31 as a potential novel therapeutic.

**Background:** DCMA is an understudied autosomal recessive disorder thought to be related to Barth syndrome but caused by mutations in *DNAJC19*, a protein of unknown function localized to the mitochondria. The clinical disease is characterized by 3-methylglutaconic aciduria, dilated cardiomyopathy, abnormal neurological development and other heterogeneous features. Until recently no effective therapies had been identified and affected patients frequently died in early childhood from intractable heart failure.

**Methods:** Dermal fibroblasts from four pediatric patients with DCMA were used to establish parameters of mitochondrial dysfunction. Mitochondrial structure, reactive oxygen species (ROS) production, cardiolipin composition and gene expression were evaluated.

**Results:** Immunocytochemistry with semi-automated quantification of mitochondrial structural metrics and transmission electron microscopy demonstrated mitochondria to be highly fragmented in DCMA fibroblasts compared to healthy control cells. Live-cell imaging demonstrated significantly increased ROS production in patient cells. These structural and functional abnormalities were reversed by treating DCMA fibroblasts with SS-31, a synthetic peptide that localizes to the inner mitochondrial membrane. Levels of cardiolipin were not significantly different between control and DCMA cells and were unaffected by SS-31 treatment.

**Conclusions:** Our results demonstrate the abnormal mitochondrial structure and function in fibroblasts from patients with DCMA and suggest that SS-31 may represent a potential therapy for this devastating disease.

## INTRODUCTION

The dilated cardiomyopathy with ataxia syndrome (DCMA), also known as 3-methylglutaconic aciduria type V, is a rare and understudied autosomal recessive disorder caused by mutations in the poorly characterized gene DNAJ Heat Shock Protein Family (Hsp40) Member C19 (*DNAJC19*) (1-4). The DNAJ family of proteins act as molecular chaperones and are defined by their J-domains which regulate the function of HSP70 chaperones (5). DNAJC19 is localized to the inner mitochondrial membrane and although some of its interacting partners have been identified (4), its precise role is unknown. DCMA was first described in the Dariusleut Hutterite population of southern Alberta who represent the largest population of patients in the world with only sporadic cases reported elsewhere (3,6,7). In the Hutterites, a genetically-isolated population that share a common European ancestry and a communal lifestyle, DCMA is caused by a single homozygous *DNAJC19* intronic pathogenic variant NG_022933.1:c.130-1G>C (rs137854888) that leads to abnormal splicing and a truncated, non-functional protein (2). DCMA is a heterogeneous disorder characterized by 3-methylglutaconic aciduria, dilated cardiomyopathy, developmental delay, neuromotor abnormalities, growth failure, prolongation of the QT interval and various other systemic features (8). End-stage heart failure leading to death in early childhood is common and, until recently, no effective therapeutic had been identified (9). However, the mechanism of disease remains unknown.

DCMA is phenotypically related to Barth syndrome (3-methylglutaconic aciduria type II) which is caused by mutations in the X-linked *TAZ* gene and whose clinical features partially overlap those seen in DCMA (10,11). *TAZ* encodes the tafazzin protein which is involved in the remodeling of cardiolipin (CL), a phospholipid predominantly localized to the inner mitochondrial membrane (11). CL has important roles in stabilizing mitochondrial membrane protein complexes and maintaining mitochondrial structure and membrane curvature (12). CL acyl chain remodelling is disrupted in cardiomyopathy, including Barth syndrome, and heart failure (13-16). In cultured cells, knock-down of *DNAJC19* expression was reported to affect CL remodeling, which may explain the related clinical features of DCMA and Barth syndrome (4). Although this *in vitro* data demonstrated that *DNAJC19* deficiency resulted in changes in CL composition and abnormal mitochondrial structure and dysfunction, results from DCMA patients have been conflicting. Both decreased and normal electron transport chain complex activities in tissues and cells have been reported (3,6,7), with Al Teneiji et al. reporting normal mitochondrial morphology in skeletal muscle (7). Despite the conflicting findings, the potential for abnormal mitochondrial structure and function in DCMA may represent a possible target for therapeutic intervention. The Szeto-Schiller peptide SS-31 (also known as elamipretide or Bendavia) interacts specifically with CL to affect membrane curvature and prevent peroxidative damage (17-19) and has shown pre-clinical promise as a treatment for mitochondrial disorders and heart failure (20-22). Our study aimed to characterize the structure and dysfunction of mitochondria found in primary dermal fibroblasts isolated from pediatric DCMA patients and to evaluate the effect of treatment with SS-31.

## MATERIALS AND METHODS

### Fibroblasts

After obtaining informed consent, clinically-indicated skin biopsies were obtained from pediatric patients undergoing investigation for metabolic disease. Fibroblasts were expanded in the Molecular Genetics Laboratory at the Alberta Children’s Hospital and subsequently frozen at −80°C until use. Four fibroblast strains from patients with biochemically and/or genetically-confirmed DCMA were selected for this study. Commercially-available control fibroblast strains derived from healthy adults or children were obtained from ThermoFisher Scientific or the Coriell Institute. All fibroblasts were grown in T25 or T75 cell culture flasks (ThermoFisher Scientific) with Minimum Essential Medium Eagle supplemented with 10% fetal bovine serum, 1 mM sodium pyruvate, 2 mM glutamine, 200 μM uridine, and 100 U/ml penicillin-streptomycin (Sigma-Aldrich). Cells were maintained under mycoplasma-free and sterile conditions in a tissue culture incubator equilibrated with 5% CO_2_ at 37°C and medium was changed every five days. SS-31 was synthesized by China Peptides (23). Experiments using SS-31 were performed by incubating fibroblasts for 24 hours with 100 nM SS-31.

### Imaging

To prepare cells for immunocytochemistry, confluent cells were dissociated using trypsin-EDTA then collected by centrifugation at 2,000 rpm for 10 min. Cell pellets were resuspended in fresh medium post-passage and seeded onto individual sterilized microscope coverslips placed on the bottom of a 24-well tissue culture plate. Cells were then allowed to grow for 48-h prior to staining. Cells on glass coverslips were washed twice with Dulbecco’s phosphate-buffered saline (DPBS) then fixed with pre-warmed 4% paraformaldehyde (J.T. Baker) in DPBS and incubated at 37°C for 15 min. Cells were then washed three times with DPBS, quenched with 50 mM NH_4_Cl for 15 min at room temperature (RT) then washed again with DPBS and stored at 4°C. When ready to stain, cells were permeabilized with 0.2% Triton X-100 in PBS for 15 min then washed three times with DPBS, blocked with 10% FBS for 25 min at RT then incubated with 1:1000 TOMM20 primary antibody (Sigma-Aldrich, cat. HPA011562) diluted in 5% FBS for 1-h at 37°C. Cells were then washed three times (5 min per wash) with 5% FBS diluted in DPBS. Cells were then incubated with the AlexaFluor 488 secondary antibody (1:1000, ThermoFisher Scientific, cat. A11034) in 5% FBS for 1-h at RT. Cells were washed then stored at 4°C in the dark until imaged on a Zeiss LSM880 confocal microscope using a 63X oil objective.

To quantify mitochondrial fragmentation in fibroblasts, thirty TOMM20-stained cells per cell line and treatment were manually graded based on a set fragmentation scale. Hyperfused cells were assigned a grade of (1), cells with intermediate fragmentation were assigned (2), and a grade of (3) was assigned to cells exhibiting substantial mitochondrial fragmentation (24). Significance was determined using a two-way ANOVA with a Holm-Sidak correction for multiple comparisons. To quantitatively assess cellular mitochondrial networks in an objective manner, a semi-automated ImageJ plug-in Mitochondrial Network Analysis (MiNA) toolset was used (25). Briefly, TOMM20-stained fibroblasts were imaged using a Zeiss LSM880 high-resolution confocal microscope. Images were then randomly cropped to select 30 individual cells. These 30 cells were identical to the those used for manual quantification. Cells were then pre-processed by using ImageJ functions unsharp mask, CLAHE, and median filtering then batch processed through MiNa. Raw data from MiNa was put through R Studio (ggbiplot, vegan, readxl, plyr, scales, and grid packages) to generate the PCA plots and calculate significant differences in clustering through Adonis tests. MiNa output displays mean network size, mean fragment length, and mitochondrial footprint. Mean network size is calculated through counting the number of mitochondrial branches per network. Mean fragment length refers to the average mitochondrial rod/branch length. Mitochondrial footprint is described as the total area in the cell expressing mitochondrial marker TOMM20. Significance was determined using a two-way ANOVA with a Holm-Sidak correction for multiple comparisons.

To assess mitochondrial ultrastructure using transmission electron microscopy, DCMA and control fibroblasts were cultured in 24-well plates to over 80% confluence. Once grown, cells were fixed and sent to University of Calgary’s Microscopy and Imaging Facility. Processed cells were imaged using a Hitachi H7650 transmission electron microscope.

### Reactive Oxygen Species (ROS) Production

Fibroblasts cultured on 35 mm glass plates (World Precision Instruments) to 50% confluence were co-stained with MitoSOX Red (ThermoFisher Scientific, cat. M36008) and MitoTracker Green (ThermoFisher Scientific, cat. M7514). Fresh fibroblast medium (2 mL) containing 5 μM MitoSOX Red and 70 nM MitoTracker Green was added to the cells and incubated for 20 min at 37°C. Cells were then washed with DPBS and new medium was added. Cells were incubated at 37°C for 20 min for de-staining and then imaged using an Olympus spinning disc confocal system (Olympus SD OSR) operated using Metamorph software. Cells were then analyzed using ImageJ. Briefly, each cell was isolated through a selection tool on both treatment images. Once identified, remaining fluorescence in the image was cleared. The cells were then subjected to a defined threshold to keep brightness consistent. Both channels were then combined using the image calculator resulting in a cell expressing co-localized fluorescence. Mean grey intensity of the cells was then calculated and plotted using GraphPad Prism 7. Seventy individual cells were quantified per cell line and treatment. Significance was determined through a one-way ANOVA with a Holm-Sidak correction for multiple comparisons.

### Western Blotting

Control and patient fibroblasts were seeded onto T25 flasks and allowed to grow overnight at 37°C and 5% CO_2_. Cells were then treated with 100 nM SS-31 or vehicle control for 24-h. Subsequently, cells were harvested, cell pellets washed and lysed with RIPA buffer containing protease inhibitors (Amersco, cat. M250). Total cell lysates (20 μg) were resolved on SDS-PAGE gels and transferred onto PVDF membranes. Blots were probed with antibodies against OPA1 (BD Bioscience, cat. 612606) at 1:1000 final dilution followed by horseradish peroxidase-conjugated secondary antibodies. Blots were finally incubated with Clarity ECL substrate (Biorad) according to manufacturer’s instructions and imaged on an Amersham Imager AI600. Densitometric analysis of band intensities were performed using ImageJ and normalized to a loading control (HSP60). Data was plotted using Prism 7 (GraphPad Software) and significance was determined using an one-way ANOVA followed by a Tukey correction for multiple comparisons.

### RNA Preparation and RNA-Seq Analysis

Total RNA was extracted from DCMA fibroblasts (n=4) and control fibroblasts (n=4) using the RNA extraction Mini kit (Invitrogen) according to the manufacturer’s protocol. RNA purity was assessed and quantified using Nanodrop and a Qubit 2.0 fluorometer (ThermoFisher Scientific). The sequencing library was prepared using 2 μg of RNA and the TruSeq Stranded mRNA library preparation kit (Illumina). RNA sequencing generating single-end 100 base pair reads was performed on the Illumina NextSeq500 platform. Raw FASTQ files were generated using Illumina NextSeq Control software (version 2.02). For RNA-Seq analysis, initial sequencing quality was inspected using FASTQC. Next, transcript counts were estimated using kallisto (26) with reference genome GRCh37 and the default settings. Kallisto-estimated counts were then summarized to the gene level using the tximport package in RStudio. Differential gene expression from the counts data was performed using the Bioconductor package DESeq2. Read counts for control and patient fibroblasts were compared to determine the log2-fold change in abundance for each transcript. Raw p-values were adjusted for multiple comparisons with the Benjamini-Hochberg method.

### Cardiolipin Analysis

CL mass and species composition were determined as previously described (27).

### Statistical Analysis

The data are presented as mean ± standard deviation (SD) and analyzed as described above. A p value < 0.05 was considered significant.

## RESULTS

### Patient characteristics

Dermal fibroblasts have often been used to study mitochondrial dysfunction in human diseases (28-30). The fibroblasts used in this study came from four individual Hutterite children from three different families with distinct clinical phenotypes (**Table 1**) despite harboring the same homozygous pathogenic variant. All patients had evidence of dilated cardiomyopathy by echocardiography with a globular and/or dilated left ventricle. Two patients (D1 and D2) had mild left ventricular dysfunction with a left ventricular ejection fraction (LVEF) of 40-50% (normal >50%) and two patients (D3 and D4) had severe dysfunction with a LVEF <35%. Each patient also had other comorbidities, most commonly developmental delay, a prolonged QT interval on the electrocardiogram and failure to thrive. The patients with severe cardiac dysfunction were both deceased at the time of this study. All studies were approved by the Conjoint Health Research Ethics Board at the University of Calgary.

**Table 1.**
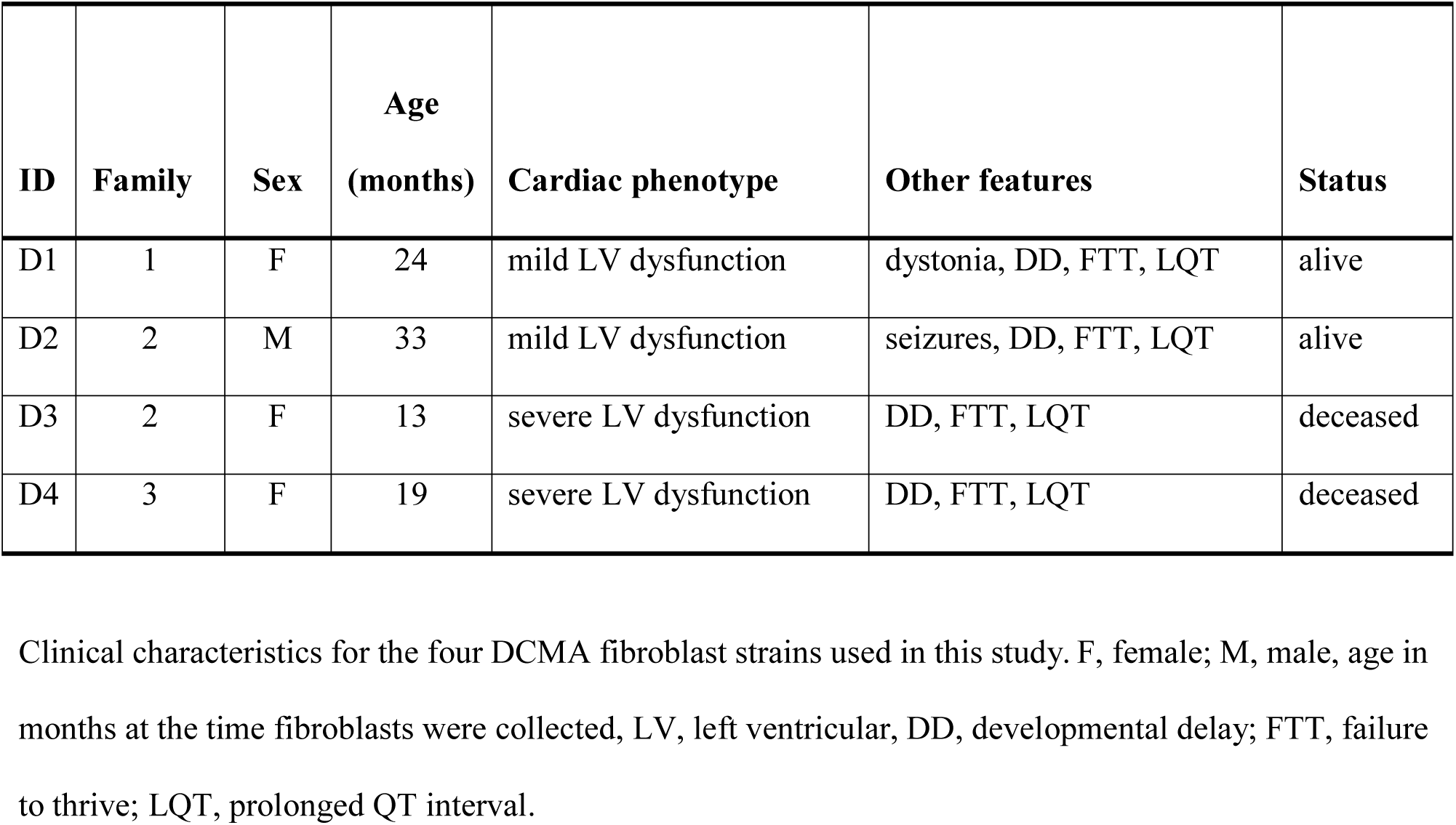
Patient information.

### Mitochondrial fragmentation in DCMA fibroblasts is reversible by incubation with SS-31

TOMM20, an outer mitochondrial membrane protein, was stained to elucidate mitochondrial structure in DCMA and control fibroblasts. Qualitatively, mitochondrial networks in all DCMA fibroblasts appeared fragmented and disorganized in contrast to control cells which displayed intact and reticular mitochondrial networks (**Figure 1A**). After 24 hours of incubation with 100 nM SS-31, the mitochondrial networks in the DCMA fibroblasts qualitatively appeared to be less fragmented and more net-like with increased branching of mitochondrial networks and longer fragments (**Figure 1B**). Semi-automated analysis of mitochondrial structure was used to quantify network size, fragment length and mitochondrial footprint (25). All parameters were found to be significantly lower in the DCMA cells in comparison to controls and the addition of SS-31 resulted in significant improvement in all three mitochondrial metrics (**Figure 1C-E**). Principal components analysis (PCA) encompassing all three mitochondrial metrics was performed for all cells in the presence and absence of SS-31 (**Figure 1F**). The DCMA patient cells clustered together and were significantly different (p < 0.0001) from the control cells. In the presence of SS-31, the DCMA cells exhibited significant improvement (p < 0.0001) in the combined mitochondrial metrics, migrating away from the untreated cells and towards the control cluster. Manual grading of mitochondrial fragmentation was performed to confirm the accuracy of our semi-automated quantification. Fibroblasts from DCMA patients displayed a higher percentage of intermediate and fragmented cells compared to control and, following treatment with SS-31, DCMA patient cell lines exhibited more hyperfused and intermediate mitochondria with a lower relative percentage of fully fragmented cells (**Supplemental Figure 1**). To further evaluate mitochondrial structure and the effect of SS-31, transmission electron microscopy (TEM) of a single DCMA strain (D1) and control fibroblast strain was performed before and after treatment with 100 nM SS-31 for 24 hours. The resulting high-magnification images showed that, qualitatively, mitochondria in the DCMA cells appeared less dark, indicating a lower electron density, and had thinner individual cristae, abnormalities that disappeared after SS-31 exposure (**Figure 2**).

**Figure 1.**
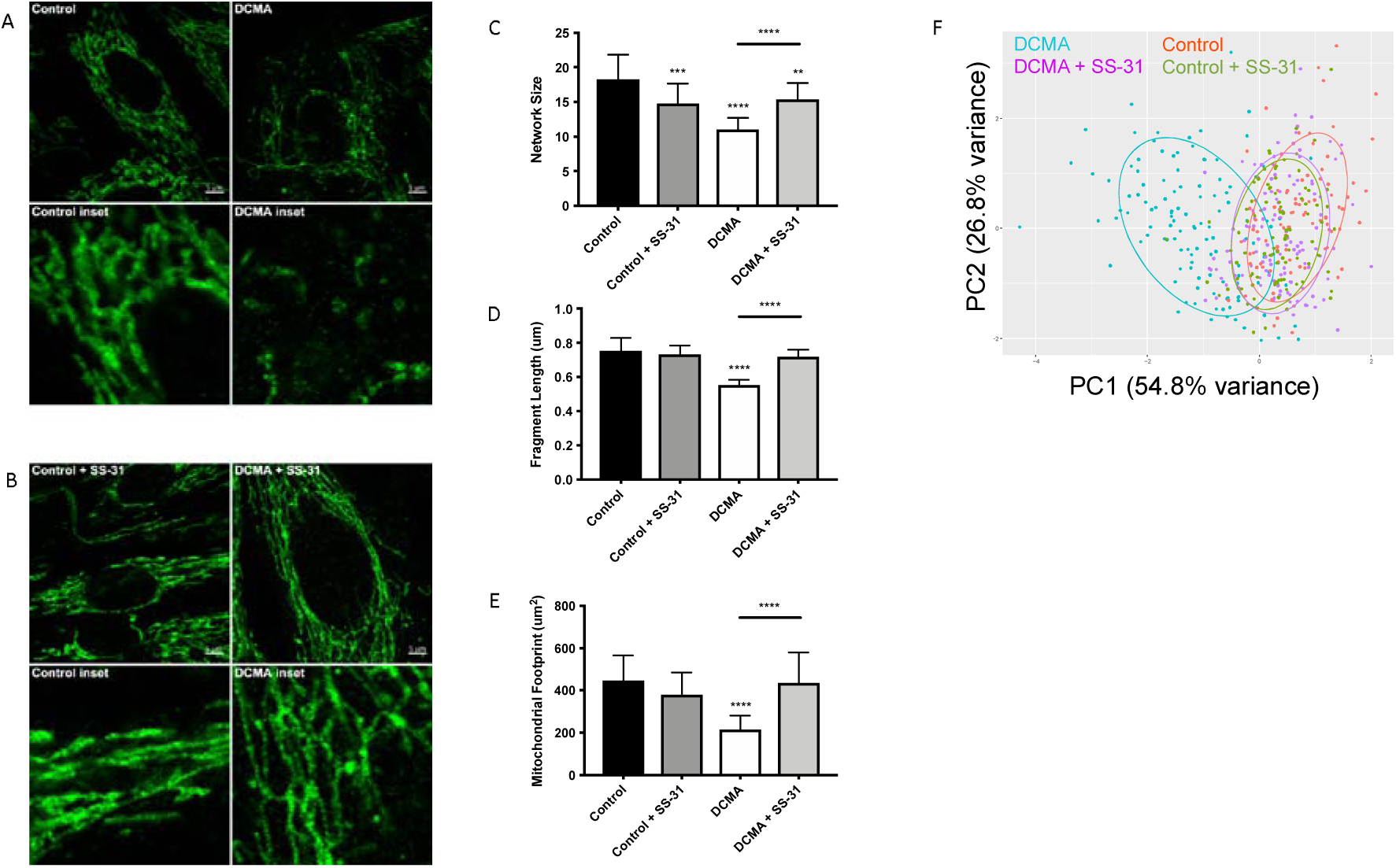
Visualization and quantification of TOMM20 staining of mitochondria. (A) Representative example of TOMM20 staining of control and DCMA fibroblasts. (B) TOMM20 staining of control and DCMA fibroblasts treated with SS-31 (100 nM for 24 hours). Scale bar measures 5 μm. Inset boxes represent the corresponding region at higher magnification. (C) Mean network size for control and DCMA fibroblasts representing the number of mitochondrial branches per network. DCMA mitochondria have significantly smaller mitochondrial networks that were restored by SS-31. (D) Mean fragment length is the average mitochondrial rod/branch length with DCMA cells having significantly smaller fragments compared to controls that increased with SS-31. (E) Mitochondrial footprint is the total area in the cell expressing mitochondrial marker TOMM20 and was significantly smaller in DCMA fibroblasts when compared to controls but increased significantly with SS-31. Data are the mean ± SD of measurements from 30 individual cells for each cell strain (n=3-4). Groups were compared using a two-way ANOVA, ** p < 0.01, *** p < 0.001, ****, p < 0.0001. (F) PCA plot incorporating data for all mitochondrial metrics (network size, fragment length and mitochondrial footprint) from control fibroblasts and DCMA fibroblasts before and after exposure to SS-31.

**Figure 2.**
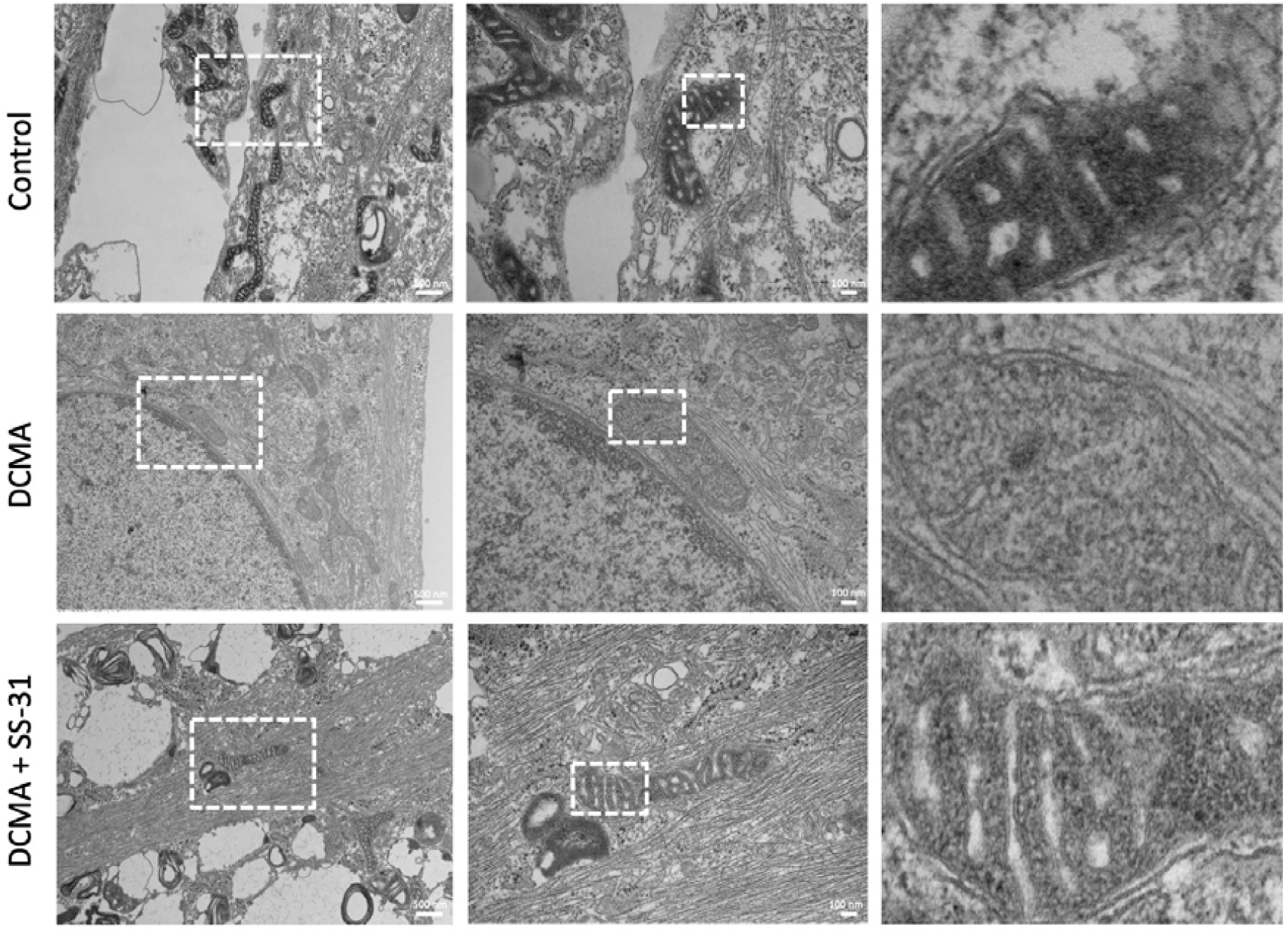
Transmission electron micrographs of mitochondria. Control and DCMA (D1) fibroblasts with and without exposure to SS-31 are shown with increasing magnification from left to right as indicated by the white dashed boxes.

### Increased ROS production in DCMA fibroblasts is reversible by incubation with SS-31

Mitochondrial ROS production was measured using live-cell imaging and specific dyes (MitoTracker Green and MitoSOX Red) to co-localize the mitochondrial network with the relative fluorescence of mitochondrial superoxide. Semi-automated mean intensity analysis of the co-localized signals showed significantly higher (p < 0.0001) mitochondrial ROS formation in the DCMA fibroblasts compared to controls. Treatment with SS-31 significantly (p < 0.0001) reduced mitochondrial ROS production in the DCMA cells and had no effect on the control cells (**Figure 3**).

**Figure 3.**
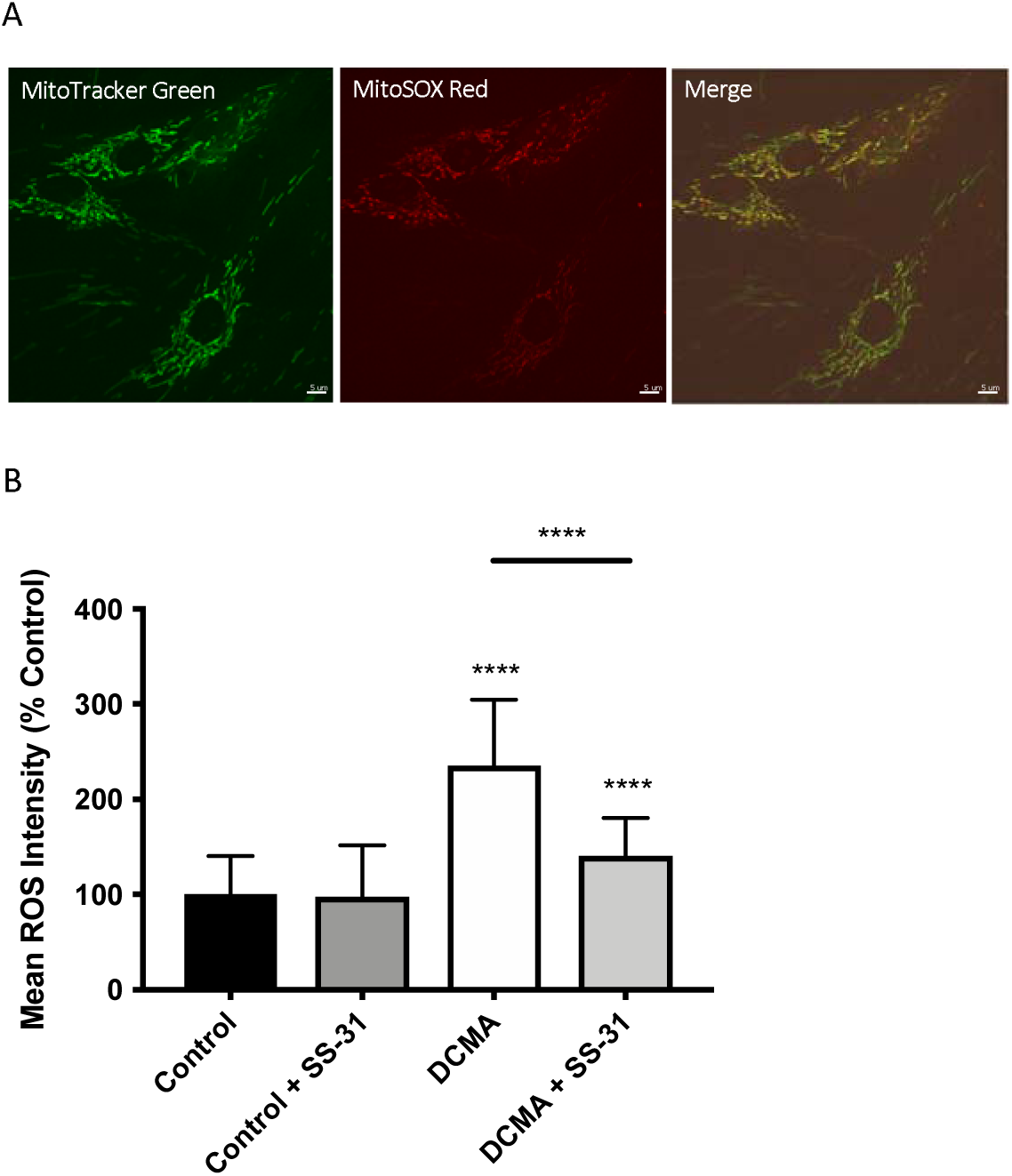
Live-cell imaging of mitochondrial ROS production. (A) Example of fibroblasts stained with MitoTracker Green, MitoSOX Red and merged images. Scale bar measures 5 μm. (B) Intensity of mitochondrial ROS staining in control and DCMA fibroblasts with and without treatment with SS-31 (100 nM for 24 hours). ROS intensity was significantly increased in DCMA fibroblasts strains but significantly decreased by SS-31. Data are mean ± SD of measurements from 70 individual cells from each control (n=1) and DCMA (n=4) fibroblast strain. Significance was determined with a one-way ANOVA and a Holm-Sidak correction for multiple comparisons. ****, p < 0.0001.

### Changes in the length of OPA1 are reversed by SS-31 in DCMA fibroblasts

Western blotting was performed on DCMA patient fibroblasts and a control to ascertain the relative ratio of the long (L-OPA1) and short (S-OPA1) isoforms of OPA1. All four DCMA patient lines showed a significant reduction in the ratio of the long and short forms that was reversed by treatment with SS-31 (**Figure 4**).

**Figure 4.**
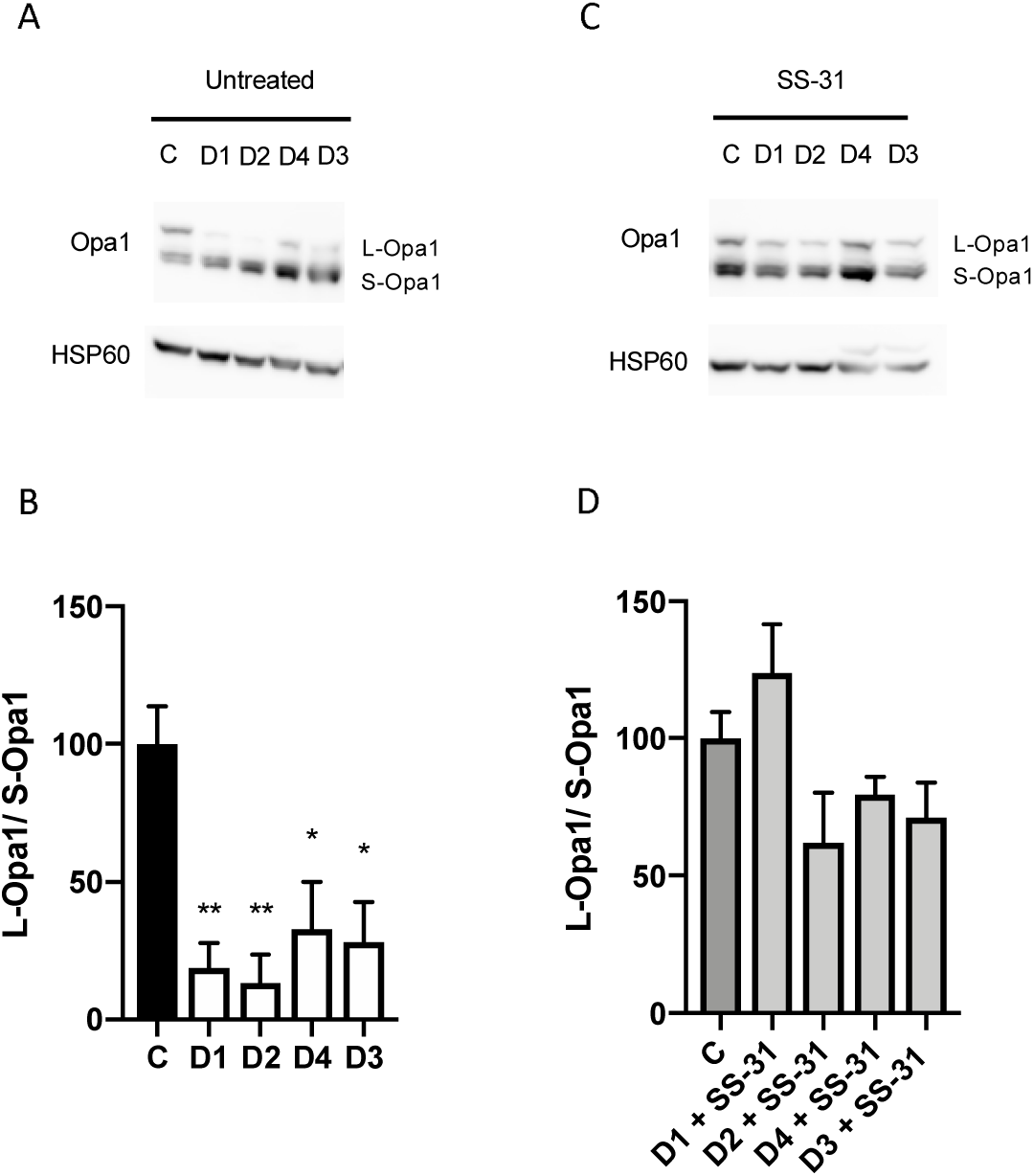
Western blotting of changes in the ratio of OPA1 isoforms. (A) Western blot of untreated control (C1) and DCMA (D1-D4) fibroblasts showing the long and short isoforms of OPA1. (B) Densitometric analysis of untreated fibroblasts with D1-D4 plotted relative to C1. The quantity of L-OPA1 is significantly reduced in DCMA cells. Data represent mean ± SD from two separate replicates. Significance was determined using a Tukey post-hoc test. *, p < 0.05; **, p ≤ 0.01. (C) Western blot of C1 and D1-D4 fibroblasts treated with 100 nM SS-31 for 24 hours showing the long and short isoforms of OPA1 and the HSP60 loading control. (D) Densitometric analysis of SS-31 treated fibroblasts. There were no significant differences between any of the groups.

### Total Cardiolipin is not Reduced in DCMA

Analysis of 22 individual molecule species of CL did not identify any significant differences between DCMA and control fibroblasts (**Supplemental Table 1**). Similarly, total CL was not significantly different between patient and control cells with or without exposure to SS-31 (**Figure 5**).

**Figure 5.**
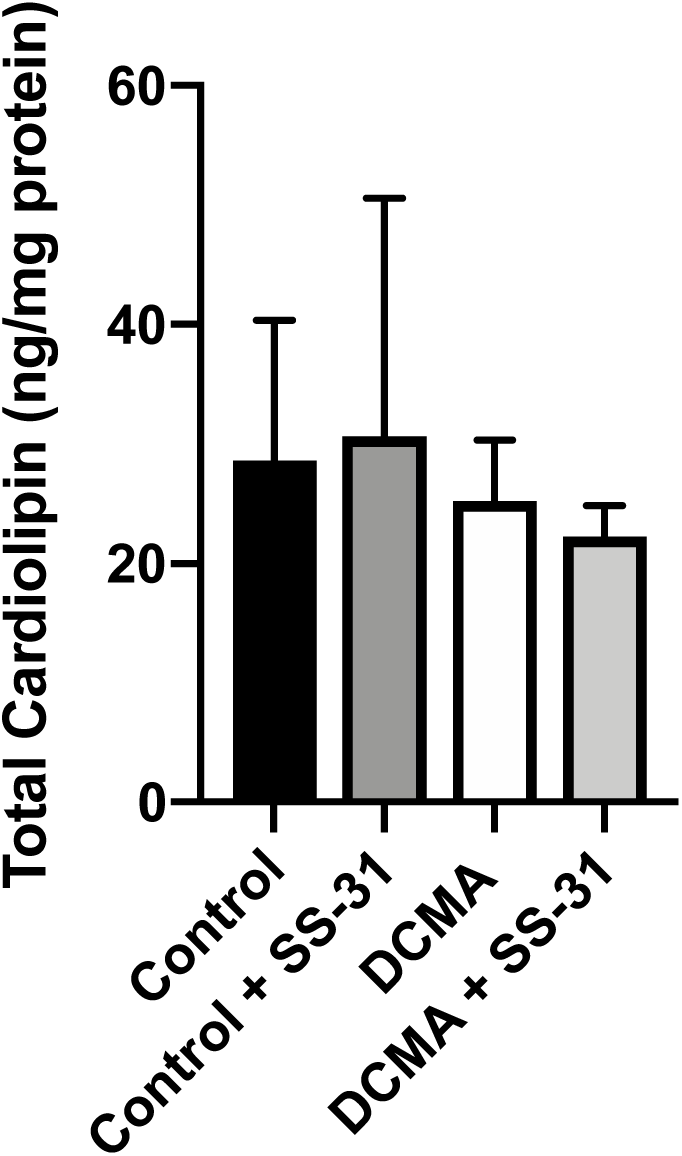
Total cardiolipin content. For control (n=3) and DCMA (n=4) fibroblasts, the total cardiolipin content was measured before and after incubation with SS-31 (100 nM for 24 hours). No significant differences were found between any of the groups.

### RNA-Seq Identifies Changes in Gene Expression Related to DCMA

Comparison between DCMA and control fibroblasts identified 262 transcripts that were significantly differentially-expressed (p < 4.9 × 10^-5^). However, there were five transcripts that were highly significantly different (p < 10^-18^) (**Table 2**). Implicated genes of particular note included *DNAJC19* and those involved in oxidative stress (*GSTM1*) and mitochondrial biogenesis (*GATD3A*) (31,32).

**Table 2.** RNA-Seq results. Most significantly differentially-expressed genes in DCMA fibroblasts.

| **ID** | **Log2-fold change** | **Adjusted P value** | **Gene Symbol** |
| --- | --- | --- | --- |
| XM_005270782.5 | 4.95 | $5.46 \times 10^{-24}$ | *GSTM1* |
| NM_145261.3 | -10.87 | $1.79 \times 10^{-23}$ | *DNAJC19* |
| XM_017028479.1 | -11.59 | $4.33 \times 10^{-23}$ | *GATD3A* |
| NM_001282418.1 | 10.21 | $1.70 \times 10^{-18}$ | *STAG2* |
| NM_005049.2 | -10.64 | $4.22 \times 10^{-18}$ | *PWP2* |

## DISCUSSION

Using dermal fibroblasts collected from four individual children with DCMA, we have identified defects in mitochondrial structure and function, specifically abnormal morphology and increased ROS production. Our results support previous *in vivo* and *in vitro* observations characterizing DCMA as a mitochondrial disease, provide a previously-lacking characterization of mitochondrial form and function in DCMA patient fibroblasts and demonstrate a striking response to the novel peptide therapeutic SS-31.

Immunocytochemistry for the outer mitochondrial membrane protein TOMM20 demonstrated that mitochondria in DCMA fibroblasts were severely fragmented with significantly reduced mitochondrial fragment length, network size and total mitochondrial footprint. TEM provided additional insight into the mitochondrial abnormalities induced by mutated DNAJC19, demonstrating that the electron density and cristae thickness were severely reduced. Reduced electron density, reflected by a decrease in the relative darkness of the mitochondrial matrix, suggests that DCMA mitochondria are likely to be in a lower energetic state in comparison to control mitochondria which are in a more condensed state and therefore actively phosphorylating ADP to produce cellular energy (33).

The abnormal mitochondrial structure in DCMA fibroblasts was associated with significantly higher ROS production. ROS are implicated in numerous roles, including cellular signalling, and in the correct balance are critically important for maintaining homeostasis and proper cellular function (34,35). An increase in ROS production can cause oxidative stress and subsequent peroxidative damage, particularly of cardiolipin which is very susceptible to this type of injury due to its composition and location (11). From our observations, it is not clear if the increased ROS production is the primary insult or secondary to the abnormal mitochondrial structure. The mitochondrial structural abnormalities that we visualized are consistent with an imbalance between mitochondrial fission and fusion. This conclusion is supported by our finding that DCMA cells exhibited significantly lower proportions of the L-OPA1 isoform which is required for mitochondrial fusion and cristae formation (36). A similar loss of L-OPA1 was observed in genetically-modified HEK293T cells and associated with abnormalities in CL composition (4). However, despite the abnormalities in mitochondrial structure and OPA1 isoform proportions, we did not see a significant difference in either total CL or individual CL species between DCMA and control fibroblasts. Given the purported link between DCMA and Barth syndrome (based on the presence of excess 3-methylglutaconic acid), this finding was unexpected. Given the documented abnormalities in CL in Barth syndrome (37), this finding suggests that the underlying cause of disease in DCMA and Barth syndrome will be different. Our RNA-Seq results support our observations of abnormal mitochondrial structure and function but do not immediately suggest mechanism.

Despite our lack in insight into disease mechanism, the mitochondrially-targeted peptide SS-31 shows promise as a therapeutic for DCMA, paralleling results seen for other mitochondrial disorders and heart failure (21,22). Incubating cells with SS-31 for just 24 hours, and using a concentration similar to that previously documented to be safe and effective *in vitro* (38), the overall mitochondrial structure in patient fibroblasts improved significantly both qualitatively and quantitatively. In addition to improved mitochondrial structure, the amount of mitochondrially-produced ROS also significantly decreased with exposure to SS-31. Although SS-31 improved mitochondrial structure and reduced oxidative stress in DCMA cells, similar to the effects observed in Friedreich ataxia (a neurodegenerative disease also associated with cardiomyopathy related to mitochondrial dysfunction), the precise mechanism of action is not known (21). However, recent work suggests that SS-31 improves coupling of electron transport chain complexes CI and CIV which may be responsible for reducing ROS production (22). We have observed reduced CI and CIV complex activity in skeletal muscle and liver from DCMA patients (Khan, unpublished data). Alternatively, due to its antioxidant activity, SS-31 may be reducing ROS abundance or it may be specifically protecting cardiolipin from peroxidative damage. Our results showing no significant changes in the levels of CL are consistent with those recently published showing that SS-31 appears to exert its effect by influencing the function of the electron transport chain rather than affecting CL directly (22). Interestingly, SS-31 significantly improved the expression of L-OPA1 and resulted in a healthier balance of L-OPA1/S-OPA1 in our DCMA cells. CL has been associated with L-OPA1 and it is hypothesized that their interaction results in adequate mitochondrial fusion (39). Through protection of CL, SS-31 could be improving the interaction of L-OPA1 with CL and may provide an explanation for the improved mitochondrial structure seen in our cells post-treatment.

Although metabolically quiescent, our research has shown dermal fibroblasts to be an adequate *in vitro* model for mitochondrial structural abnormalities. As such, the effect of other potential therapeutics could be evaluated using our fibroblasts. For example, the cardiac glycoside digoxin has recently been shown to improve myocardial function and structure in children with DCMA but the impact of digoxin on mitochondrial structure and function remains to be evaluated (9).

In conclusion, we have completed a novel *in vitro* study of the rare mitochondrial disease DCMA using patient-derived dermal fibroblasts. Analysis of mitochondrial morphology identified multiple abnormalities of mitochondrial structure that may be contributing to elevated ROS production and decreased organelle fusion. The observation that SS-31 is able to ameliorate all of these abnormalities is also a novel and exciting finding. Since dysfunctional mitochondria most likely underlie the lethal cardiomyopathy frequently found in this disorder, identification of SS-31 as a potential therapeutic may have important future clinical implications.

## Abbreviations List

CL: cardiolipin
DCMA: dilated cardiomyopathy with ataxia syndrome
DD: developmental delay
FTT: failure to thrive
LQT: prolonged QT interval
LV: left ventricular
LVEF: left ventricular ejection fraction
PCA: principal components analysis
ROS: reactive oxygen species
TEM: transmission electron microscopy

## Acknowledgements

We would like to thank Vincent Ebacher in the Hotchkiss Brain Institute for his image analysis support. We also acknowledge the imaging resources of the Charbonneau Microscopy Facility and the Microscopy and Imaging Facility at the University of Calgary.

## Author contributions

PM, XW, AK, TS and SCG conceived and designed the experiments. PM, XW, TZ, RS, MK and AR performed experiments and acquired data. PM, JH and FI performed data analysis. BA and DS provided reagents. PM, XW and SCG wrote the manuscript. All authors read and approved the manuscript.

---

**Supplemental Figure 1.**
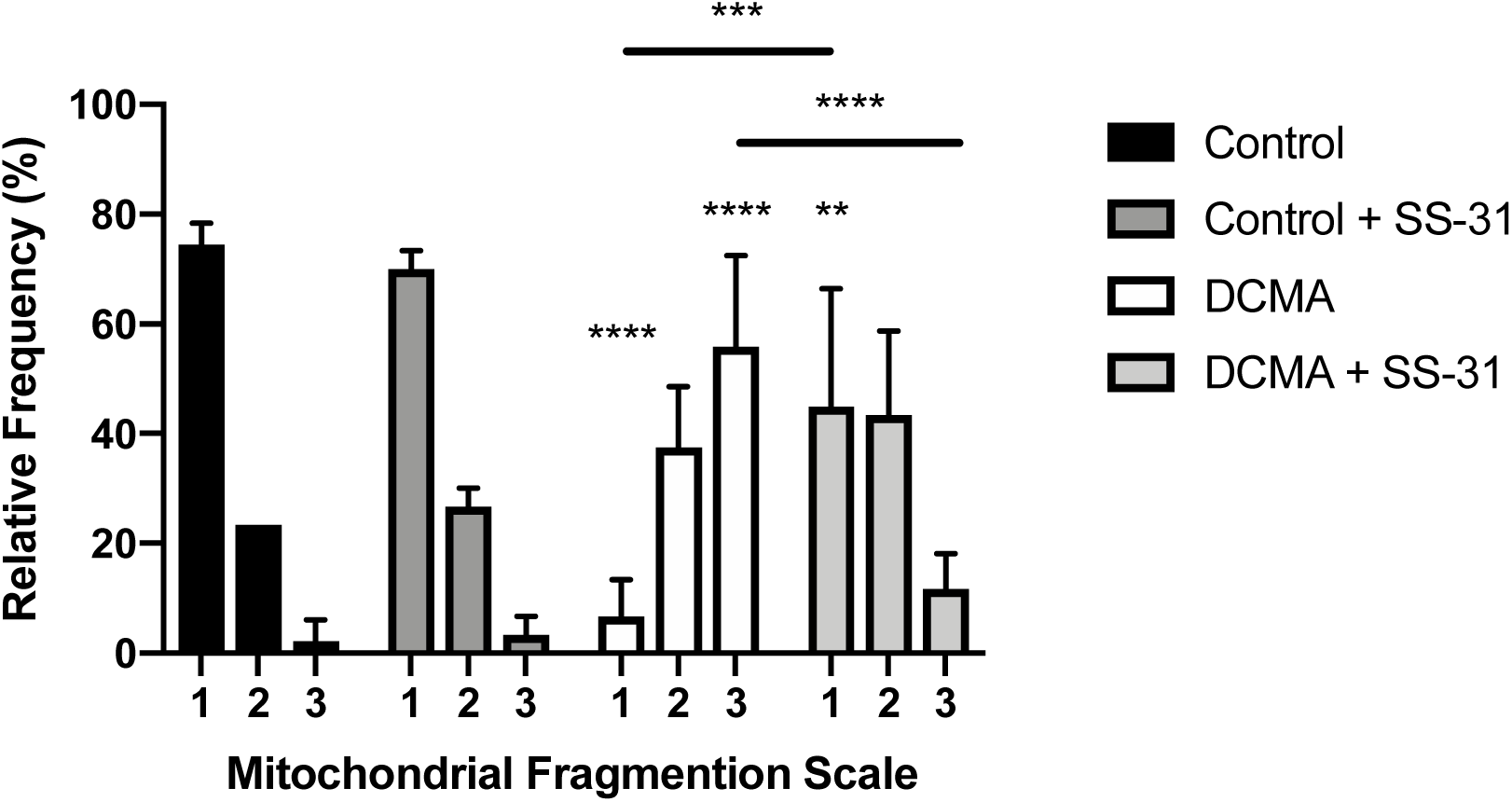
Manual quantification of mitochondrial morphology. Mitochondrial morphology from 30 individual cells for control (n=3) and DCMA (n=4) fibroblasts before and after treatment with SS-31 (100 nM for 24 hours) was quantified according to a three-point fragmentation scale: (1) hyperfused, (2) intermediate and (3) fragmented. Data represent mean ± SD. Significance was determined using a two-way ANOVA with a Holm-Sidak correction for multiple comparisons. ** p < 0.01, *** p < 0.001, **** p < 0.0001.

**Supplemental Table 1.** Cardiolipin species in control and DCMA fibroblasts before and after exposure to SS-31. Data are mean ± SD for control (n=3) and DCMA (n=4) cell strains. No significant differences (p < 0.05) were detected between the groups by Student’s t-test.

| Species | Control | Control + SS-31 | DCMA | DCMA + SS-31 |
| --- | --- | --- | --- | --- |
| CL (18:1)(18:2)<sub>2</sub>(20:1) | 0.36 ± 0.15 | 0.35 ± 0.23 | 0.28 ± 0.16 | 0.27 ± 0.1 |
| CL (18:1)(18:2)<sub>2</sub>(20:1); CL(18:1)<sub>2</sub>(18:2)(20:2) | 0.46 ± 0.22 | 0.37 ± 0.22 | 0.29 ± 0.1 | 0.36 ± 0.08 |
| CL (18:1)(18:2)<sub>2</sub>(20:2) | 1.75 ± 0.89 | 2.01 ± 1.46 | 1.42 ± 0.53 | 1.34 ± 0.31 |
| CL (18:1)(18:2)<sub>2</sub>(22:6) | 0.10 ± 0.04 | 0.13 ± 0.12 | 0.07 ± 0.03 | 0.07 ± 0.05 |
| CL (18:1)<sub>2</sub>(18:2)(20:1); CL(18:1)<sub>2</sub>(18:0)(20:3) | 0.19 ± 0.13 | 0.18 ± 0.13 | 0.17 ± 0.07 | 0.15 ± 0.05 |
| CL (18:1)<sub>2</sub>(18:2)(20:2) | 0.81 ± 0.49 | 0.82 ± 0.53 | 0.72 ± 0.27 | 0.74 ± 0.22 |
| CL (18:2)(18:0)(22:6)<sub>2</sub>; CL(18:1)<sub>2</sub>(22:6)<sub>2</sub> | 0.08 ± 0.03 | 0.05 ± 0.02 | 0.07 ± 0.04 | 0.09 ± 0.03 |
| CL (18:2)(18:1)(20:3)(22:6) | 0.09 ± 0.04 | 0.06 ± 0.03 | 0.09 ± 0.03 | 0.10 ± 0.08 |
| CL (18:2)(18:1)<sub>3</sub> | 9.20 ± 4.11 | 10.30 ± 7.47 | 9.20 ± 1.98 | 7.85 ± 1.47 |
| CL (18:2)<sub>2</sub>(18:1)(16:1) | 2.71 ± 1.08 | 2.84 ± 2.31 | 2.19 ± 0.42 | 1.49 ± 0.95 |
| CL (18:2)<sub>2</sub>(18:1)(20:3); CL(18:2)<sub>3</sub>(20:2) | 0.50 ± 0.3 | 0.58 ± 0.37 | 0.39 ± 0.12 | 0.39 ± 0.05 |
| CL (18:2)<sub>2</sub>(18:1)(22:5) | 0.10 ± 0.05 | 0.09 ± 0.06 | 0.05 ± 0.01 | 0.07 ± 0.02 |
| CL (18:2)<sub>2</sub>(18:1)(22:6) | 0.36 ± 0.2 | 0.48 ± 0.44 | 0.20 ± 0.07 | 0.19 ± 0.04 |
| CL (18:2)<sub>3</sub>(16:1) | 1.51 ± 0.34 | 1.69 ± 1.12 | 1.28 ± 0.47 | 1.21 ± 0.18 |
| CL (18:2)<sub>3</sub>(18:1) | 5.13 ± 1.89 | 5.20 ± 2.94 | 4.71 ± 1.01 | 4.02 ± 1.02 |
| CL (18:2)<sub>3</sub>(20:3) | 0.64 ± 0.4 | 0.68 ± 0.44 | 0.42 ± 0.14 | 0.47 ± 0.04 |
| CL (18:2)<sub>3</sub>(20:4) | 0.50 ± 0.31 | 0.47 ± 0.28 | 0.27 ± 0.12 | 0.22 ± 0.15 |
| CL (18:2)<sub>3</sub>(22:5) | 0.10 ± 0.05 | 0.15 ± 0.12 | 0.05 ± 0.02 | 0.05 ± 0.01 |
| CL (18:2)<sub>4</sub> | 1.29 ± 0.4 | 1.41 ± 0.66 | 1.08 ± 0.51 | 1.07 ± 0.13 |
| CL (18:3)(18:2)<sub>3</sub> | 0.20 ± 0.09 | 0.24 ± 0.16 | 0.15 ± 0.04 | 0.10 ± 0.07 |
| CL (C18:2)<sub>2</sub>(C18:1)<sub>2</sub> | 2.47 ± 1.06 | 2.45 ± 1.56 | 1.99 ± 0.65 | 1.94 ± 0.21 |
| LysoCL (18:2)<sub>3</sub> | 0.11 ± 0.05 | 0.10 ± 0.04 | 0.11 ± 0.07 | 0.08 ± 0.03 |

## REFERENCES

1. Blomen VA, Majek P, Jae LT et al. Gene essentiality and synthetic lethality in haploid human cells. Science 2015;350:1092–1096.

2. Davey KM, Parboosingh JS, McLeod DR et al. Mutation of DNAJC19, a human homologue of yeast inner mitochondrial membrane co-chaperones, causes DCMA syndrome, a novel autosomal recessive Barth syndrome-like condition. J Med Genet 2006;43:385–393.

3. Ojala T, Polinati P, Manninen T et al. New mutation of mitochondrial DNAJC19 causing dilated and noncompaction cardiomyopathy, anemia, ataxia, and male genital anomalies. Pediatr Res 2012;72:432–7.

4. Richter-Dennerlein R, Korwitz A, Haag M et al. DNAJC19, a mitochondrial cochaperone associated with cardiomyopathy, forms a complex with prohibitins to regulate cardiolipin remodeling. Cell Metab 2014;20:158–71.

5. Kampinga HH, Craig EA. The HSP70 chaperone machinery: J proteins as drivers of functional specificity. Nat Rev Mol Cell Biol 2010;11:579–92.

6. Ucar SK, Mayr JA, Feichtinger RG, Canda E, Coker M, Wortmann SB. Previously Unreported Biallelic Mutation in DNAJC19: Are Sensorineural Hearing Loss and Basal Ganglia Lesions Additional Features of Dilated Cardiomyopathy and Ataxia (DCMA) Syndrome? JIMD Rep 2016:8.

7. Al Teneiji A, Siriwardena K, George K, Mital S, Mercimek-Mahmutoglu S. Progressive Cerebellar Atrophy and a Novel Homozygous Pathogenic DNAJC19 Variant as a Cause of Dilated Cardiomyopathy Ataxia Syndrome. Pediatr Neurol 2016;62:58–61.

8. Sparkes R, Patton D, Bernier F. Cardiac features of a novel autosomal recessive dilated cardiomyopathic syndrome due to defective importation of mitochondrial protein. Cardiol Young 2007;17:215–7.

9. Greenway SC, Dallaire F, Hazari H, Patel D, Khan A. Addition of Digoxin Improves Cardiac Function in Children With the Dilated Cardiomyopathy With Ataxia Syndrome: A Mitochondrial Cardiomyopathy. Can J Cardiol 2018;34:972–977.

10. Clarke SL, Bowron A, Gonzalez IL et al. Barth syndrome. Orphanet J Rare Dis 2013;8:23.

11. Dudek J, Maack C. Barth syndrome cardiomyopathy. Cardiovasc Res 2017;113:399–410.

12. Osman C, Voelker DR, Langer T. Making heads or tails of phospholipids in mitochondria. J Cell Biol 2011;192:7–16.

13. Chicco AJ, Sparagna GC. Role of cardiolipin alterations in mitochondrial dysfunction and disease. Am J Physiol Cell Physiol 2007;292:C33–44.

14. Saini-Chohan HK, Holmes MG, Chicco AJ et al. Cardiolipin biosynthesis and remodeling enzymes are altered during development of heart failure. J Lipid Res 2009;50:1600–8.

15. Sparagna GC, Chicco AJ, Murphy RC et al. Loss of cardiac tetralinoleoyl cardiolipin in human and experimental heart failure. J Lipid Res 2007;48:1559–70.

16. Acehan D, Xu Y, Stokes DL, Schlame M. Comparison of lymphoblast mitochondria from normal subjects and patients with Barth syndrome using electron microscopic tomography. Laboratory investigation; a journal of technical methods and pathology 2007;87:40–48.

17. Szeto HH. First-in-class cardiolipin-protective compound as a therapeutic agent to restore mitochondrial bioenergetics. Br J Pharmacol 2014;171:2029–50.

18. Szeto HH, Birk AV. Serendipity and the discovery of novel compounds that restore mitochondrial plasticity. Clin Pharmacol Ther 2014;96:672–83.

19. Szeto HH, Schiller PW. Novel therapies targeting inner mitochondrial membrane--from discovery to clinical development. Pharm Res 2011;28:2669–79.

20. Birk AV, Liu S, Soong Y et al. The mitochondrial-targeted compound SS-31 re-energizes ischemic mitochondria by interacting with cardiolipin. Journal of the American Society of Nephrology: JASN 2013;24:1250–61.

21. Zhao H, Li H, Hao S et al. Peptide SS-31 upregulates frataxin expression and improves the quality of mitochondria: implications in the treatment of Friedreich ataxia. Sci Rep 2017;7:9840.

22. Chatfield KC, Sparagna GC, Chau S et al. Elamipretide Improves Mitochondrial Function in the Failing Human Heart. JACC Basic to translational science 2019;4:147–157.

23. Wu J, Hao S, Sun XR et al. Elamipretide (SS-31) Ameliorates Isoflurane-Induced Long-Term Impairments of Mitochondrial Morphogenesis and Cognition in Developing Rats. Front Cell Neurosci 2017;11:119.

24. Sabouny R, Fraunberger E, Geoffrion M et al. The Keap1-Nrf2 Stress Response Pathway Promotes Mitochondrial Hyperfusion Through Degradation of the Mitochondrial Fission Protein Drp1. Antioxidants & redox signaling 2017;27:1447–1459.

25. Valente AJ, Maddalena LA, Robb EL, Moradi F, Stuart JA. A simple ImageJ macro tool for analyzing mitochondrial network morphology in mammalian cell culture. Acta histochemica 2017;119:315–326.

26. Bray NL, Pimentel H, Melsted P, Pachter L. Near-optimal probabilistic RNA-seq quantification. Nat Biotechnol 2016;34:525–7.

27. Ravandi A, Leibundgut G, Hung MY et al. Release and capture of bioactive oxidized phospholipids and oxidized cholesteryl esters during percutaneous coronary and peripheral arterial interventions in humans. J Am Coll Cardiol 2014;63:1961–71.

28. James AM, Wei YH, Pang CY, Murphy MP. Altered mitochondrial function in fibroblasts containing MELAS or MERRF mitochondrial DNA mutations. The Biochemical journal 1996;318 (Pt 2):401–7.

29. Onesto E, Colombrita C, Gumina V et al. Gene-specific mitochondria dysfunctions in human TARDBP and C9ORF72 fibroblasts. Acta neuropathologica communications 2016;4:47.

30. Kogot-Levin A, Saada A, Leibowitz G et al. Upregulation of Mitochondrial Content in Cytochrome c Oxidase Deficient Fibroblasts. PLoS One 2016;11:e0165417.

31. Masuda T, Wada Y, Kawamura S. ES1 is a mitochondrial enlarging factor contributing to form mega-mitochondria in zebrafish cones. Sci Rep 2016;6:22360.

32. Ponamarev MV, She YM, Zhang L, Robinson BH. Proteomics of bovine mitochondrial RNA-binding proteins: HES1/KNP-I is a new mitochondrial resident protein. Journal of proteome research 2005;4:43–52.

33. Hackenbrock CR. Ultrastructural bases for metabolically linked mechanical activity in mitochondria. II. Electron transport-linked ultrastructural transformations in mitochondria. J Cell Biol 1968;37:345–69.

34. Tsutsui H, Kinugawa S, Matsushima S. Mitochondrial oxidative stress and dysfunction in myocardial remodelling. Cardiovasc Res 2009;81:449–56.

35. Ray PD, Huang BW, Tsuji Y. Reactive oxygen species (ROS) homeostasis and redox regulation in cellular signaling. Cellular signalling 2012;24:981–90.

36. Olichon A, Guillou E, Delettre C et al. Mitochondrial dynamics and disease, OPA1. Biochim Biophys Acta 2006;1763:500–9.

37. Mejia EM, Zinko JC, Hauff KD, Xu FY, Ravandi A, Hatch GM. Glucose Uptake and Triacylglycerol Synthesis Are Increased in Barth Syndrome Lymphoblasts. Lipids 2017;52:161–165.

38. Hao S, Ji J, Zhao H et al. Mitochondrion-Targeted Peptide SS-31 Inhibited Oxidized Low-Density Lipoproteins-Induced Foam Cell Formation through both ROS Scavenging and Inhibition of Cholesterol Influx in RAW264.7 Cells. Molecules 2015;20:21287–97.

39. Liu R, Chan DC. OPA1 and cardiolipin team up for mitochondrial fusion. Nature cell biology 2017;19:760–762.

